# Molecular autism research in Africa: a scoping review comparing publication outputs to Brazil, India, the UK, and the USA

**DOI:** 10.1101/2022.11.11.516128

**Authors:** Emma Frickel, Sophia Bam, Erin Buchanan, Caitlyn Mahony, Mignon van der Watt, Colleen O’Ryan

**Affiliations:** Department of Molecular and Cell Biology, University of Cape Town, Private Bag, Rondebosch, 7701 Cape Town, South Africa

**Keywords:** Autism, autism spectrum disorder, genetics, molecular, Sub-Saharan Africa, South Africa, Brazil, India, UK, USA

## Abstract

The increased awareness of autism spectrum disorders (ASD) is accompanied by burgeoning ASD research, and concerted research efforts are trying to elucidate the molecular ASD aetiology. However, much of this research is concentrated in the Global North, with recent reviews of research in Sub-Saharan Africa (SSA) highlighting the significant shortage of ASD publications from this region. The most limited focus area was molecular research with only two molecular studies ever published from SSA, both being from South Africa (SA). We examine the molecular ASD research publications from 2016 to 2021 from all African countries, with a special focus on SA. The SSA publications are compared to Brazil and India, two non-African, low-to-middle-income countries (LMICs), and to the UK and USA, two high-income countries (HICs). There were 228 publications across all regions of interest; only three publications were from SA. Brazil (n=29) and India (n=27) had almost 10 times more publications than SA. The HICs had more publications than the LMICs, with the UK (n=62) and the USA (n=74) having approximately 20 to 25 times more publications than SA, respectively. Given that SA has substantial research capacity as demonstrated by its recent research on SARS-CoV-2, we explore potential reasons for this deficit in molecular ASD publications from SA. We compare mental health research outputs, GDP per capita, research and development expenditure, and the number of psychiatrists and child psychiatrists per 100,000 people across all regions. The UK and the USA had significantly higher numbers for all these indicators, consistent with their higher publication output. Among the LMICs, SA can potentially produce more molecular ASD research, however, there are numerous barriers that need to be addressed to facilitate increased research capacity. These include cultural stigmas, challenges in accessing mental healthcare, shortages of specialists in the public sector, and the unreliability of ASD diagnostic tools across the 11 official SA languages. The unique genetic architecture of African populations presents an untapped reservoir for finding novel genetic loci associated with ASD. Therefore, addressing the disparity in molecular ASD research between the Global North and SSA is integral to global advancements in ASD research.

## INTRODUCTION

Autism spectrum disorders (ASD) are a leading cause of disability in children and adolescents (Divan *et al*., 2021). The global prevalence of lifelong neurodevelopmental disorders is currently estimated to be 1 in 100 children and is continually increasing (Zeidan *et al*., 2022). Although ASD research has increased over the years, the majority of this research has been done in high-income countries (HICs). Little is known about ASD in low-to-middle-income countries (LMICs), despite 95% of children with ASD and other developmental disabilities living in these settings (de Vries, 2016; Olusanya *et al*., 2018). Sub-Saharan Africa (SSA) comprises mostly low-income countries and has a population of more than 1 billion (World Bank, 2021). There have been no population-based prevalence studies of ASD in SSA and only 0.5% of the world’s ASD research has been done in this region (Franz *et al*., 2017). There is an urgent need to address this research disparity, especially considering the prediction that 37% of all children on the globe will live in Africa by 2050 (UNICEF, 2014).

The heritability of ASD is undisputed, but the specific genetic and molecular mechanisms involved remain elusive (Lord *et al*., 2020). As these mechanisms have crucial implications for treatment interventions, concerted research efforts aim to understand the molecular mechanisms of ASD. The underrepresentation of African populations is not limited to ASD but has been widely documented in all genetic disease studies (Oni-Orisan *et al*., 2021; Sirugo *et al*., 2019). For instance, only 3% of genome-wide association studies (GWAS) have included African populations, whereas 81% have been undertaken in European populations (Popejoy & Fullerton 2016; Fatumo *et al*., 2022). This disparity is detrimental since genetic associations may not be applicable across populations given differences in genetic structure (Oni-Orisan *et al*., 2021; Sirugo *et al*., 2019). Moreover, the well-documented, rich genetic diversity of African populations represents a unique resource with the potential to make valuable contributions to global ASD research (Campbell & Tishkoff, 2008; Sirugo *et al*., 2019). In addition to the opportunity for discovering novel genetic variants associated with ASD, the reduced linkage disequilibrium in African genomes increases the power for genetic fine-mapping (Campbell & Tishkoff, 2008; Sirugo *et al*., 2019; Tishkoff & Verrelli, 2003). This could yield novel insights into molecular mechanisms involved in ASD aetiology.

The considerable shortage of ASD research in Africa, particularly molecular studies, was evident in the three most recent reviews of ASD research in African regions (Abubakar *et al*., 2016; Bakare *et al*., 2022; Franz *et al*., 2017). Abubakar *et al*., (2016) reported the volume and scope of ASD research from Sub-Saharan Africa (SSA) over 80 years up to June 2016. The authors identified 47 publications from SSA, most of which (n=25) were from South Africa (SA). Franz *et al*., (2017) reviewed all ASD research ever published in SSA up to October 2015 and identified 53 SSA publications, 28 from SA. They also found that SSA had published 96 times fewer articles than North America over the same period (Franz *et al*., 2017). Bakare *et al*., (2022) identified 41 studies in a review of all ASD research in West Africa from 1978 to March 2021. Together, the three reviews highlight the dearth of ASD research in these regions compared to other continents. Furthermore, molecular ASD studies were the most limited of the study types, with only four such publications identified across all three reviews, two from SA and two from Nigeria. This illustrates a significant shortage of molecular studies from SSA and highlights a major knowledge gap in African ASD research. Expanding molecular research to include understudied populations is essential to develop a field that is inclusive and internationally relevant. Moreover, this is a particularly promising avenue for future research due to the vast genetic heterogeneity of autism.

Despite the limited study output from SSA, Abubakar *et al*., (2016) reported a steady increase in publications over the decade leading up to 2016. Given this promising trajectory, this scoping review has three objectives.

Firstly, to provide an update on the SA ASD publication volume and scope from 2016 to 2021, since most of the prior SSA publications were from SA. Secondly, this review focused on molecular ASD research and examined the publications from all African countries in comparison to two non-African LMICs (Brazil and India), and two HICs (the UK and the

USA). Thirdly, this review addresses the challenges that impact the landscape of ASD research in the SA context and the unique contributions that SA and African studies can make to ASD research on a global scale. This data could be used to inform future ASD research in SA and SSA.

## METHODS

### Search Strategy

This scoping review was completed using the Arksey and O’Malley framework (Arksey & O’Malley, 2005; Levac *et al*., 2010). Briefly, we identified the research question, captured the studies for screening, selected the relevant studies for inclusion, charted the data, and finally, we collated, summarised and reported our results.

We conducted searches of PubMed and Scopus, and we searched AfricaWide, CINAHL, PsychArticles and PsychInfo through EBSCOhost, to identify peer-reviewed original research publications from 1 January 2016 to 31 December 2021. The search strategy used three sets of search terms, combined in each database with “AND”, to identify ASD molecular research in the respective geographical regions of interest. The first set of search terms was “Autism Spectrum Disorder” OR “autism” OR “ASD” OR “autistic” OR “Autistic Disorder”. The second set of search terms used a combination of keywords such as “genetic” OR “molecular”. The third set specified the region, i.e., “South Africa” OR “South African”, and six separate searches were done for each region: SA, Africa, Brazil, India, the UK, and the USA. We included a search that was not limited to molecular research to capture the volume and scope of all ASD publications from SA between 2016 and 2021. This was done using the first set of search terms combined with “South Africa” OR “South African”, using “AND”.

All research articles were exported to an EndNote 20 library keeping each geographical region separate and then uploaded to Rayyan (Ouzzani *et al*., 2016), which was used to remove duplicates and carry out screening for article inclusion. All publication titles and abstracts were independently reviewed by two authors, and any conflicts were resolved by a joint review of the full text. When an agreement was not reached, the senior author also reviewed the title and abstract to reach a consensus.

### Inclusion and Exclusion Criteria

Articles from the molecular ASD searches were screened for eligibility based on the title and abstract using the following inclusion criteria: i) ASD was the primary focus of the study, ii) the study was carried out within the region of interest by authors from that region, iii) the study generated primary data, and iv) the study used molecular methods or reported molecular data. An exception for the latter was made in the broader search of SA, which aimed to capture all areas of ASD research. Searches were filtered for the date of publication (1 January 2016 - 31 December 2021).

Studies that did not meet the above criteria were excluded. Dissertations and conference reports/abstracts were also excluded. Where there were numerous authors from multiple regions, studies were excluded if the main author and/or the ethics approval was not from the region of interest.

## RESULTS AND DISCUSSION

### Molecular ASD Publications From All Regions of Interest

We retrieved 2,456 articles across all regions of interest, published between 1 January 2016 and 31 December 2021 from the database searches for molecular ASD studies. Duplicate articles and articles that did not meet our eligibility criteria were removed, and a final 228 publications were included in the review (Figure 1). SA had the lowest publication output (n=3). The USA had the highest number of publications (n=74; ~25 times more than SA), followed by the UK (n=62; ~21 times more than SA), Brazil (n=29; ~10 times more than SA), and India (n=27; 9 times more than SA). There were 33 molecular ASD publications from seven of the 54 countries in Africa. North Africa contributed most of the publications: 20 from Egypt and four from Tunisia. There were nine publications from SSA: Nigeria (n=3), South Africa (n=3), Cameroon (n=1), Seychelles (n=1) and Uganda (n=1) (Figure 2).

**Figure 1:**
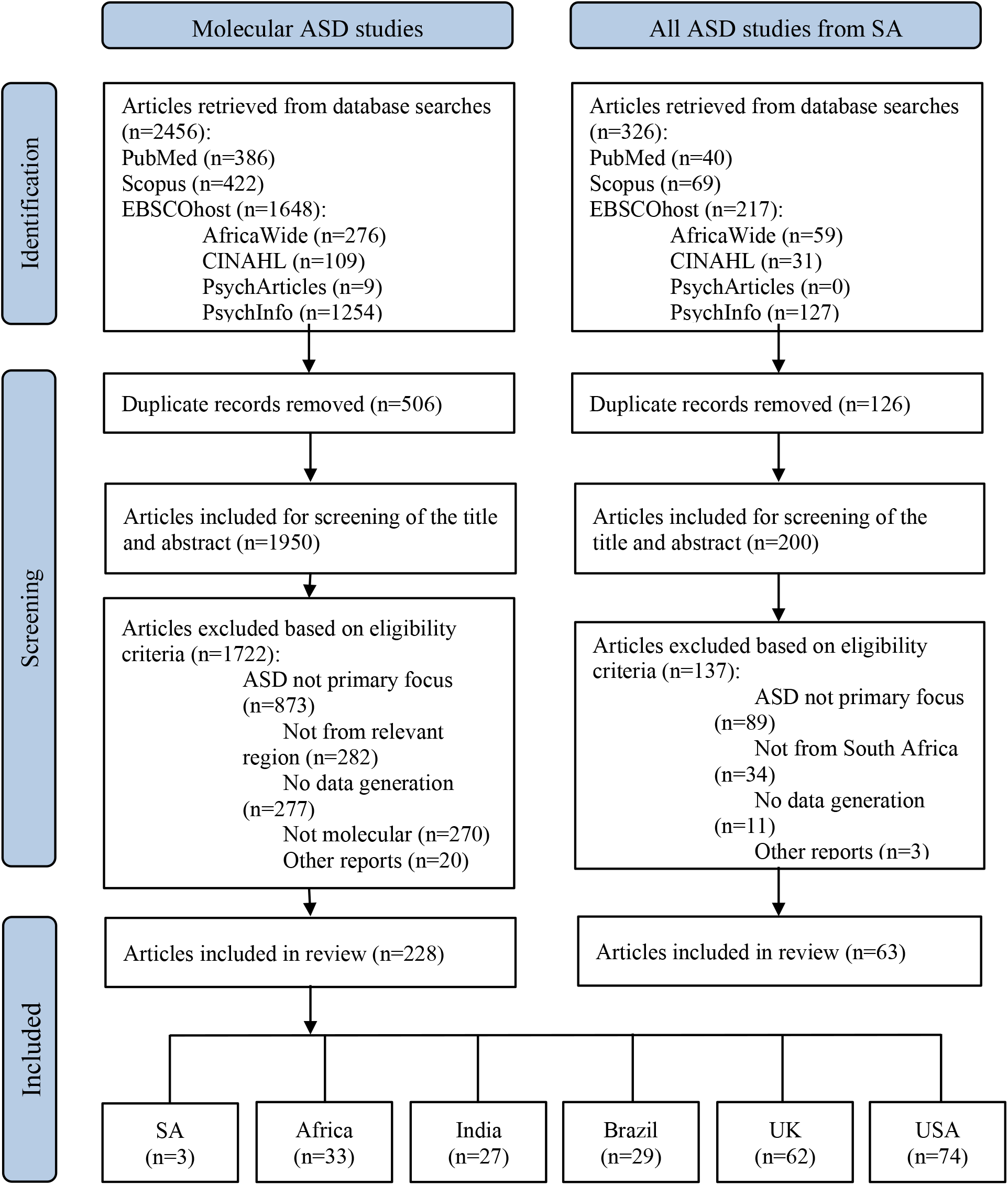
Flowchart of the study selection process for the molecular ASD studies from all geographical regions of interest (left), and for all the ASD studies from South Africa (right). The number of studies from Africa (n=33) includes the 3 from SA.

**Figure 2:**
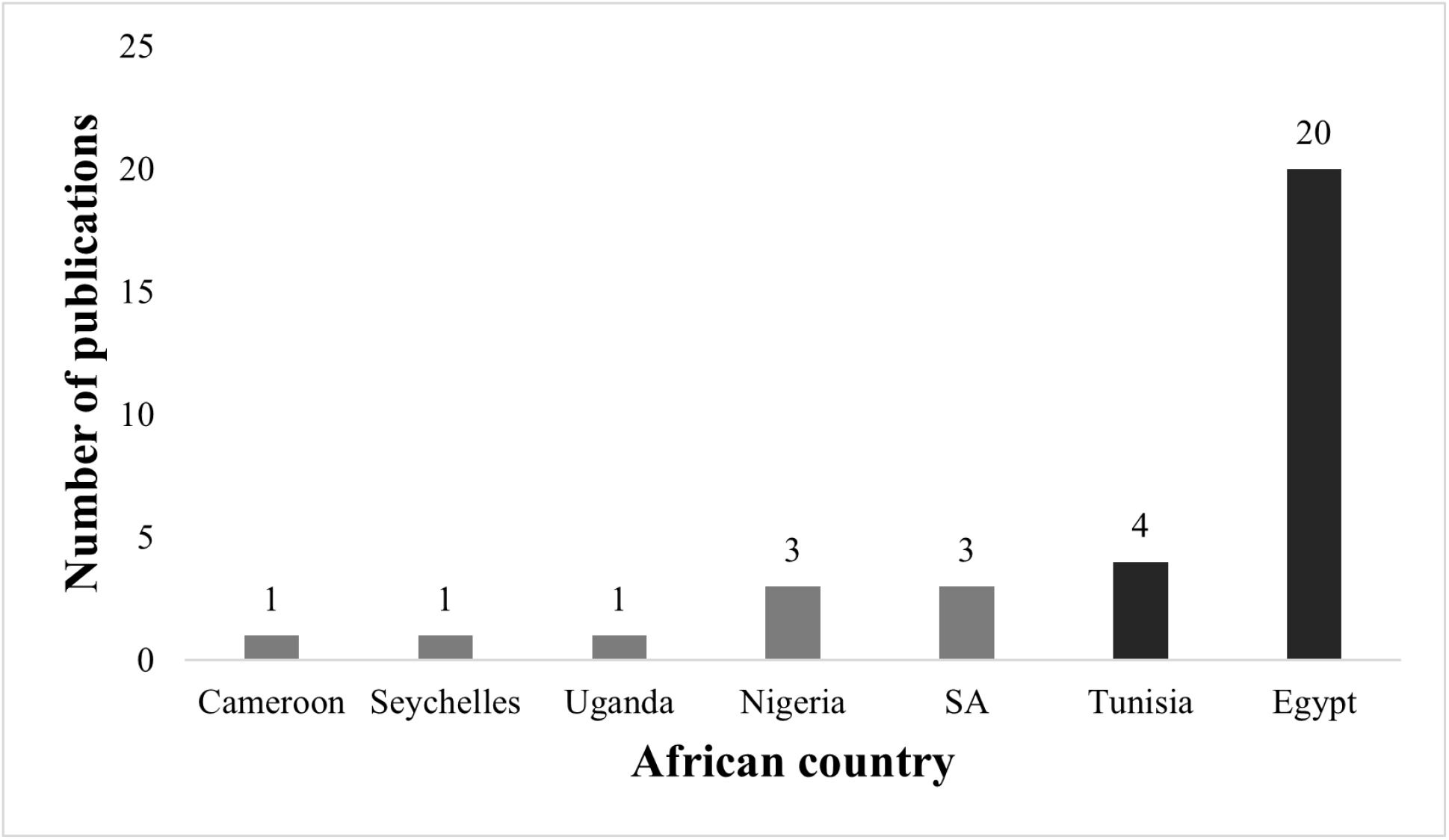
Number of molecular ASD publications from SSA countries (grey) and North African countries (black) identified in this review from 2016 to 2021.

With respect to previous reviews on ASD research outputs in these regions, we identified four reviews reporting on studies from Egypt, Nigeria, SA, and India, but were unable to find similar reports for the other regions. Egypt’s publication output far outweighed that of the six other African countries, each with less than five publications. This data is consistent with a review reporting that Egypt (n=16) produced more autism publications than all Arab countries, aside from Saudi Arabia (n=23), between 1992 and 2012 (Hussein & Taha, 2013). Three previous reviews identified four molecular publications from SSA up to 2016 from SA (n=2), Nigeria (n=1), and Tanzania (n=1) (Abubakar *et al*., 2016; Franz *et al*., 2017; Bakare *et al*., 2022). Therefore, we showed that there has been an increase in SSA autism publications from 2016 to 2021 (n=9). Given that three SA publications were identified after 2016, SA did not substantially increase their publication output, but still had the second highest number of publications from SSA. Bakare *et al*., (2022) identified two publications from Nigeria published after 2016 (Blaurock-Busch & Nwokolo Chijioke, 2018; Omotosho *et al*., 2018) that were not captured in our search, meaning that a total of five Nigerian publications after 2016 were identified. These two studies were not identified in our review due to different search databases. Bakare *et al*., (2022) searched Google, Google Scholar, and African Index Medicus, which were not included in our study. As only one publication from Nigeria was identified before 2016, Nigeria showed more of an increase in its publication output compared to the other SSA regions. This is concordant with previous reviews highlighting Nigeria as a leading contributor to ASD research in SSA. Before 2016, no studies were identified from Cameroon, Uganda, or Seychelles. The recent publications from these regions indicate an emerging interest in the field in other SSA countries and present an opportunity for collaborative research efforts. A recent scoping review of all ASD studies from India up to 15 February 2019 identified 27 genetic and biochemical (i.e., molecular) studies, 15 were published in the six years prior to 2016 (Patra & Kar., 2020). Given that we identified 27 publications (2016 - 2021), our findings show an increasing trend in publications from India in the past six years.

### Molecular ASD Publications From SA

Our search of molecular studies from SA since 2016 captured three publications, two of which were from our research group (Table 1). These two studies reported on the epigenetics of a South African ASD cohort compared to age- and sex-matched controls. The first paper described an epigenome-wide DNA methylation screen that revealed differentially methylated genes in ASD, converging on mitochondrial canonical pathways (Stathopoulos *et al*., 2020). These findings, along with urinary metabolite analysis, indicated that mitochondrial dysfunction was present in the South African ASD cohort (Stathopoulos *et al*., 2020). The second paper (Bam *et al*., 2021) examined the methylation of key mitochondrial genes involved in mitochondrial biogenesis, fission, and fusion in the same ASD cohort compared to controls. Differential methylation of these genes was correlated with increased mitochondrial DNA (mtDNA) copy number in the ASD group, indicating dysregulated mitochondrial function (Bam, *et al*., 2021). Together, the data from these two publications are consistent with previous research highlighting the role of mitochondrial dysfunction in the aetiology of ASD (Frye, 2020; Rose *et al*., 2018). The third publication from a SA research group used publicly available genetic data to characterise common and unique gene associations, and enriched pathways, in ASD, schizophrenia (SCZ), bipolar disorder (BD), and obsessive-compulsive disorder (OCD) (O’Connell *et al*., 2018). This study identified 10 common genes across the four disorders, whilst a large number of previous genetic associations were found to be disorder-specific. Commonly associated pathways included dopaminergic- and serotonergic-pathways and the hippo-signalling pathway, supporting a key role in neural development and neuronal maintenance in these disorders. The publicly available gene lists used were not specifically associated with SA individuals, therefore only two of the identified studies that generated primary data were done in SA. However, the two primary studies had a relatively small sample size, highlighting the need for large-scale genetic studies of ASD in SA, especially considering the unique genetic architecture of these populations (Campbell & Tishkoff, 2008; Sirugo *et al*., 2019).

**Table 1:** A summary of molecular ASD publications from SA between 2016 and 2021 identified in this review.

| First author | Year | Sample description | Aim of study | Summary of findings |
| --- | --- | --- | --- | --- |
| O'Connell | 2018 | Previously published gene lists for ASD, schizophrenia (SCZ), bipolar disorder (BD), and obsessive-compulsive disorder (OCD). | Characterise common and unique molecular architecture of ASD, SCZ, BD, and OCD. | 10 genes were commonly associated with ASD, SCZ, BD, and OCD, and enriched for biological pathways such as dopaminergic and serotonergic pathways. A large number of genes were disorder-specific, highlighting pathways and biological processes specific to each disorder. |
| Stathopoulos | 2020 | Buccal samples (for DNA extraction) collected from 145 SA children (93 ASD and 52 controls: African-, European-, and Mixed-ancestry). Urine (for metabolite analysis) was collected from 21 ASD children and 13 controls. | Determine epigenome-wide methylation levels to identify differentially methylated genes between ASD and control children. Assess levels of urinary metabolites between cases and controls. | 898 genes were significantly differentially methylated between ASD cases and controls. These genes converged on mitochondrial canonical pathways, implicating mitochondrial metabolism and function in ASD. Mitochondrial dysfunction was confirmed in the ASD cohort from urinary metabolite analysis. |
| Bam | 2021 | DNA was extracted from buccal samples collected from 145 SA children (93 ASD and 52 controls: African-, European-, and Mixed-ancestry). Previously published urinary metabolomic data generated from 21 ASD children and 13 controls was used (Stathopoulos *et al* 2020). | Measure DNA methylation of 6 key mitochondrial genes i.e., *PGC-1α*, *STOML*, *MFN2*, *FIS1*, *OPA1*, and *GABPA* in ASD children and controls. Additionally, determine mitochondrial DNA (mtDNA) copy number in the cohort, and examine the association with urinary metabolites. | Differential methylation of the 6 key mitochondrial genes was evident between ASD cases and controls. MtDNA copy number was elevated in ASD children and was significantly correlated with *PGC-1α* promoter methylation, and with urinary organic acids involved in mitochondrial dysfunction. These results highlight a role of mitochondrial dysfunction in ASD. |

### All Categories of ASD Research From SA

We identified 63 publications across all thematic study areas of ASD research in SA (2016 - 2021). The publication selection process is shown in Figure 1, and the thematic groupings are summarised in Figure 3. Eight focus areas emerged from the studies, namely, family and social aspects (n=25), clinical/phenotyping/behavioural studies (n=9), education (n=8), diagnosis (n=6), prevalence/demographics (n=4), professional knowledge (n=4), interventions and treatment (n=4), and molecular studies (n=3). Our data show that most of the ASD research in SA focused on family and social aspects of ASD. This is in accordance with Abubakar *et al*., (2016), but in contrast to Franz *et al*., (2017), who reported that publications on the prevalence of ASD were the most abundant. The thematic area with the lowest output across all three reviews was molecular research. Franz *et al*., (2017) found equally low numbers (n=2) in the “screening and diagnosis” category.

**Figure 3:**
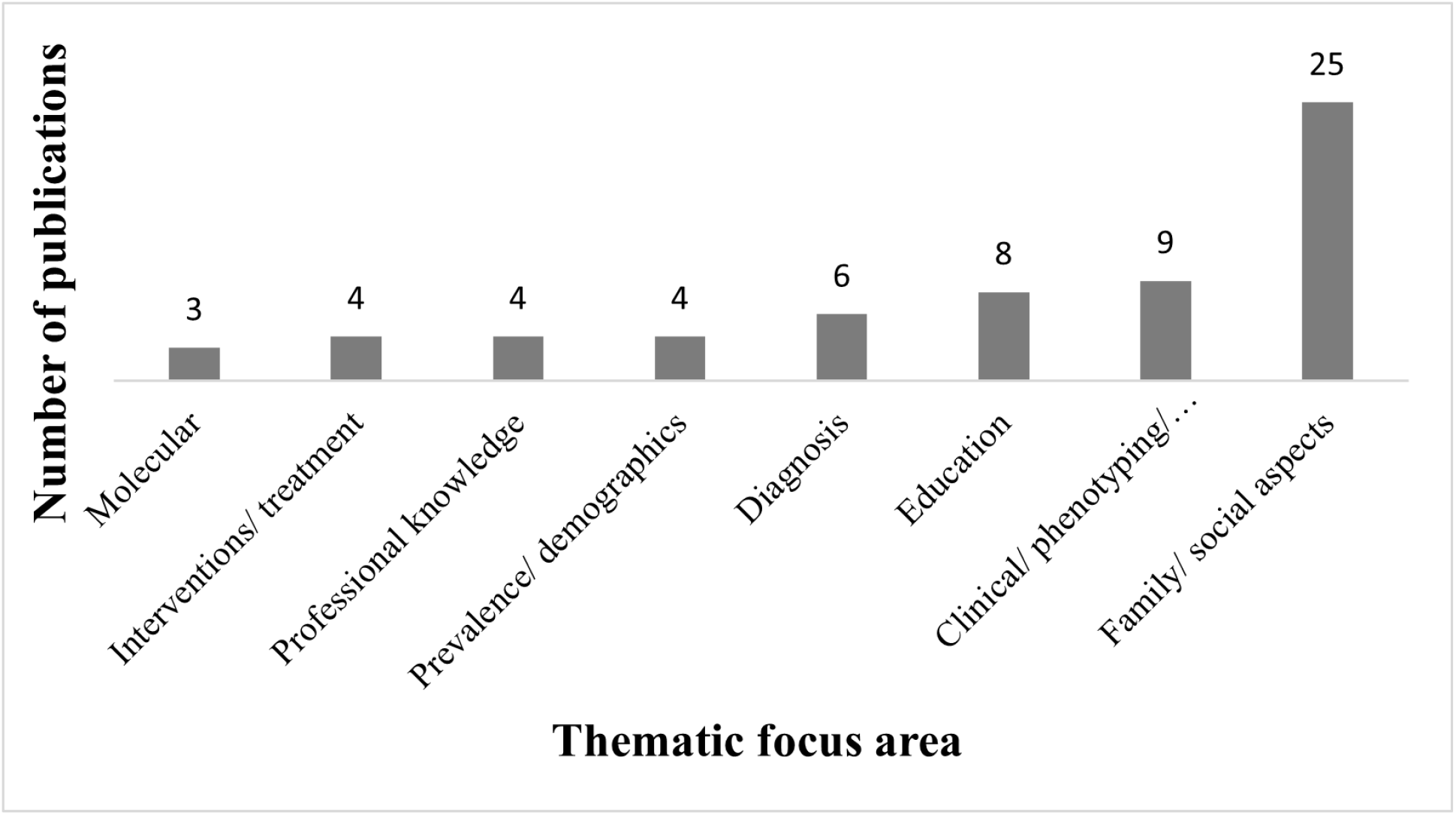
Number of South African ASD publications per thematic focus area from 2016 to 2021 identified in this review.

Franz *et al*., (2017) identified 28 ASD publications from SA up to 2015 and Abubakar *et al*., (2016) identified 26 SA ASD publications (1935 - 2016). After cross-referencing and combining their results with ours, the data showed 97 ASD studies ever published in SA up to 2021. Interestingly, 65 of these studies (67%) were published between 2016 and 2021, indicating an exponential increase in ASD research (Figure 4). However, this trend was not observed for molecular studies, since only two studies were published in SA prior to 2016 and only three studies were published after 2016. This emphasises the continued scarcity of molecular ASD research compared to other study areas.

**Figure 4:**
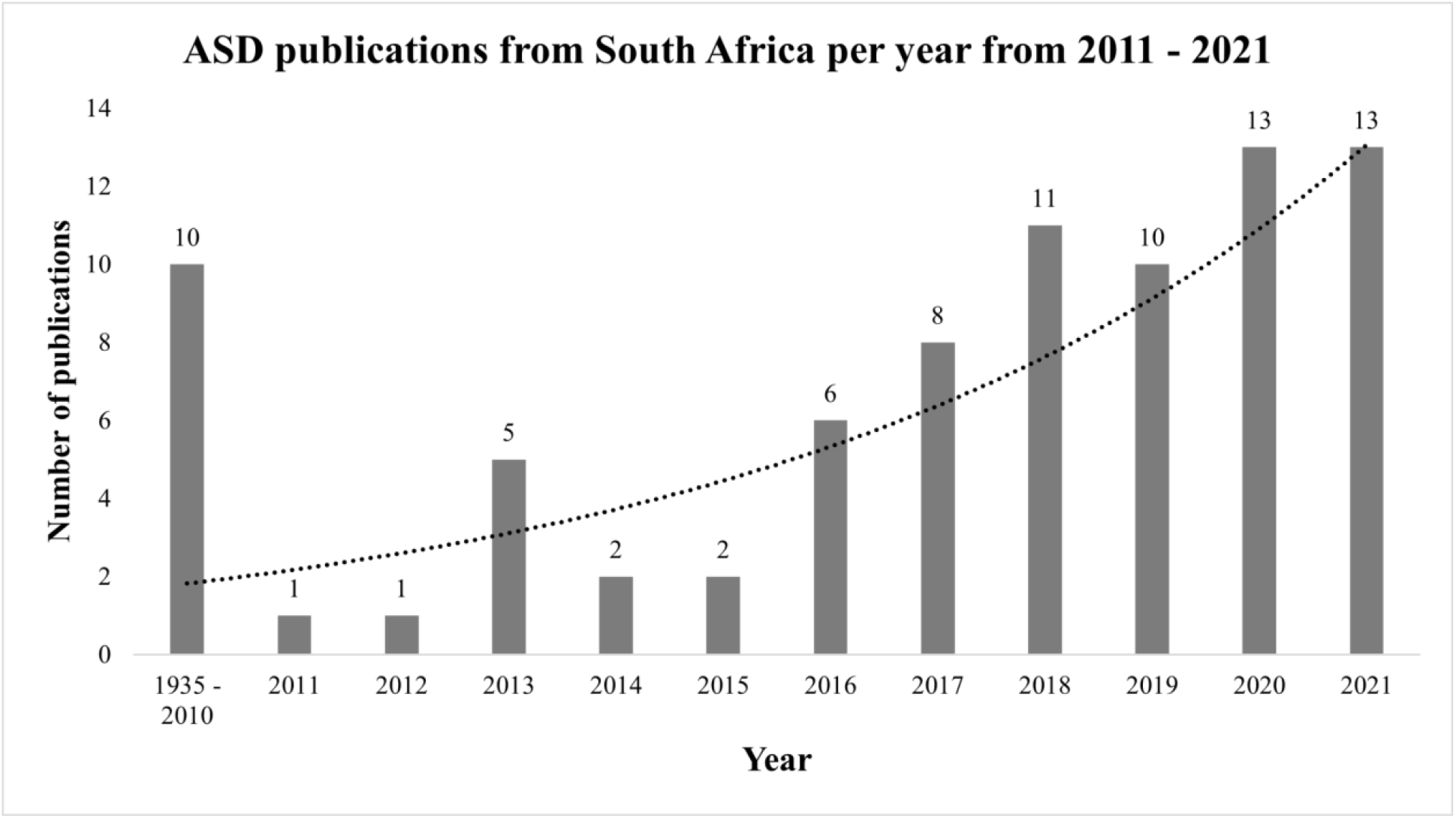
Total ASD publications from South Africa identified between 1935 and 2010 (the earliest study was published in 1970), and publications per year from 2011 – 2021. Data from 1935 to 2015 are taken from Abubakar et al. (2016) and Franz et al. (2017). Data from 2016 are from Abubakar et al. (2016) combined with our findings. Publications from 2017 onward were identified in our review.

### Mental Health Research and Resources in Regions of Interest

SA and Nigeria are the leading SSA countries in ASD research (Abubakar *et al*., 2016; Franz *et al*., 2017). Despite this, the SA publication output for molecular ASD research lags behind the other regions we examined by a substantial amount. We compared mental health research outputs to determine whether this unbalanced publication volume between regions is specific to molecular ASD research, or whether this trend is also reflected in general mental health research (World Health Organization [WHO], 2021) (Table 2). We then explored factors that may impact mental health research outputs, namely, mental health workforce expressed as the number of psychiatrists and child psychiatrists per 100,000 people, GDP per capita, and the research and development (R&D) expenditure as a percentage of GDP.

**Table 2:** Mental health research outputs, economic indicators, and mental health workforce for each region.

| Measure/ indicator |  | LMIC |  |  | HIC |  |
| --- | --- | --- | --- | --- | --- | --- |
|  |  | SA | Brazil | India | UK | USA |
| Molecular ASD publications (2016 - 2021)<sup>a</sup> |  | 3 | 29 | 27 | 62 | 74 |
| Molecular ASD publications (2019)<sup>a</sup> |  | 0 | 5 | 4 | 12 | 17 |
| Mental health publications (2019)<sup>b</sup> |  | 165 | 419 | - | 1282 | 4856 |
| Mental health research output (% of total research output; 2019)<sup>b</sup> |  | 6.89 | 5.37 | - | 12.20 | 11.45 |
| GDP per capita<sup>c</sup> | 2018 | 7,005 | 9,151 | 1,998 | 43,647 | 62,805 |
|  | 2019 | 6,625 | 8,876 | 2,072 | 43,071 | 65,095 |
|  | 2021 | 6,994 | 7,519 | 2,277 | 47,334 | 69,288 |
| Research and development expenditure (% GDP; 2018)<sup>c</sup> |  | 0.69 | 1.17 | 0.66 | 1.70 | 3.00 |
| Number of psychiatrists per 100,000 people |  | 1.59 (2019)<sup>b</sup> | 3.69 (2019)<sup>b</sup> | 0.29 (2017)<sup>d</sup> | 13.76 (2019)<sup>b</sup> | - |
| Number of child psychiatrists per 100,000 people |  | 0.11 (2019)<sup>b</sup> | - | 0.00 (2017)<sup>d</sup> | 10.31 (2019)<sup>b</sup> | - |

R&D = Research and development. Dashes (-) indicate missing data. <sup>a</sup>Data from current review; <sup>b</sup>Data for 2019, taken from The World Health Organization Mental Health Atlas 2020, Member State Profiles; <sup>c</sup>Data taken from The World Bank; <sup>d</sup>Data taken from WHO Mental Health Atlas 2017, Member State Profile (no Member State profile for India published in the WHO Mental Health Atlas 2020)

The USA and the UK had the highest number of mental health publications (29 and 8 times more than SA respectively), and compared to SA, had almost double the mental health research output as a percentage of total research output. There was no data for the mental health workforce in the USA, but the UK had the highest number of psychiatrists and child psychiatrists per 100,000 people, approximately 10 and 94 times more than SA respectively (WHO, 2020). The USA and the UK had the highest GDP per capita and R&D expenditure in 2018 (the most recent data available), substantially higher than SA (WHO, 2021). These indicators justify the higher molecular ASD research output from the UK and the USA compared to SA, as these HICs evidently have more resources available for mental healthcare and mental health research.

Considering the LMICs, Brazil had the highest number of mental health publications and psychiatrists, and the highest GDP and R&D expenditure. These indicators could partly explain the increased capacity for molecular ASD research in Brazil compared to SA. However, Brazil had more than double the mental health publications compared to SA, yet 10 times more molecular ASD publications. This shows that the disparity in molecular autism research between Brazil and SA is greater than the disparity in other mental health research. The number of mental health publications from India was not available, but there were fewer psychiatrists and child psychiatrists per 100,000 people compared to SA (Table 2). India’s GDP was 3.5 times less than SA and the difference in R&D expenditure between these regions was negligible (Table 2). Despite this, India had approximately the same number of molecular ASD publications as Brazil, nine times more than SA. Together, these data suggest that mental health research in SA is concentrated on areas outside of molecular ASD research and SA may have the capacity to channel resources and focus on ASD in the future. However, there are notable barriers specific to SA that need to be addressed in order to facilitate increased ASD research capacity.

### Unique Challenges and Opportunities for Autism Research in the SA Context

Since the emergence of the SARS-CoV-2 as a global pandemic, scientists from SA have been at the forefront of genomic surveillance allowing for the identification of 16 novel variants and real-time epidemiological analysis of Covid-19 outbreaks in SA (Tegally *et al*., 2021). This demonstrates considerable molecular research capacity in SA, yet this is not reflected in molecular autism research outputs. The shortage of molecular ASD research stems from a number of challenges culminating in limited accessibility to study participants and insufficient funding for research. Recruiting participants for ASD studies is complicated by substantial under-identification of children with autism and a lack of appropriate healthcare and educational facilities.

Receiving an ASD diagnosis in SA can be challenging due to a number of factors. There is a lack of autism awareness coupled with pervasive stigmatization in many SA communities. For example, autistic children and their families often face discrimination due to culture-bound beliefs that there are supernatural causes of autism, such as witchcraft, curses, or evil spirits (Booysen *et al*., 2021; Gona *et al*., 2015; de Vries, 2016). As a result, families often consult with traditional healers and are reluctant to seek the necessary mental healthcare (Booysen *et al*., 2021).

For those seeking help from mental healthcare services, there are further significant challenges to accessing the required support. For example, people residing in rural areas (52% of the SA population) need to travel long distances to access healthcare, with concomitant transport costs (Reid, 2006; Vergunst, 2018). Additionally, about 80% of psychiatrists in SA work in the private sector, and thus rural areas have an extremely limited number of mental health specialists (0.03 per 100,000 people) (Van Rensburg *et al*., 2022). This means that patients primarily rely on general practitioners and nurses for mental health issues. Unfortunately, access to these healthcare workers is limited due to staff shortages at treatment facilities (Booysen *et al*., 2021; Vergunst, 2018). Furthermore, there is insufficient ongoing mental health training and support for public healthcare nurses, which limits their capacity to provide adequate interventions (Petersen *et al*., 2009).

Autism diagnosis is further complicated as there are 11 official SA languages, which means that standardised diagnostic tools are not always reliable and culturally appropriate (de Vries, 2016; Durkin *et al*., 2015). This necessitates translation, adaptation, and validation of autism screening tests across all SA languages. However, many of the widely used tools have licensing restrictions and thus require permissions and payment in order to translate them into other languages which is problematic in low resource settings (Durkin *et al*., 2015). Although there have been several preliminary studies assessing the use of diagnostic instruments in some SA languages, this process is still in its infancy (Chambers *et al*., 2017; Malcolm-Smith *et al*., 2013; Smith, Malcolm-Smith & de Vries, 2017; Vorster *et al*., 2021). Furthermore, once appropriate diagnostic assessments have been established, considerable efforts will be required to adequately train professionals and effectively implement these tools (de Vries, 2016; Durkin *et al*., 2015).

The average waiting period for a child to receive a clinical diagnosis in a specialist clinic is 18 months (Guler *et al*., 2018). Following diagnosis, the next challenge is receiving the necessary educational support. SA has only nine specialised schools for autism with an estimated waiting period of more than three years for enrolment (Franz *et al*., 2018). Furthermore, the number of children on the waiting list increased by 276% between 2012 and 2016, showing an escalation in the number of children who cannot access the required educational support (Franz *et al*., 2018; Pillay *et al*., 2017).

Even if the challenges of autism diagnosis were overcome, an immense hindrance to ASD research is a lack of sufficient funding. This is largely due to the prioritisation of the infectious disease crisis. Africa is tremendously overburdened with communicable diseases, e.g., human immunodeficiency virus and acquired immunodeficiency syndrome (HIV/AIDS), tuberculosis (TB) and malaria (Boutayeb, 2010; Bhutta *et al*., 2014). In 2022, Africa had 25.6 million HIV cases, the highest of all WHO regions, and at 7.5 million, SA had the most HIV cases of all countries globally (WHO, 2022). The research capacity for HIV, TB, malaria and more recently SARS-CoV-2, has been facilitated by increased foreign (e.g., European Commission) and local funding (Arvanitis *et al*., 2022; National Health Research Committee, 2022). Consequently, this has created significant setbacks for mental health research in Africa (Bakare et al., 2014; de Vries, 2016). For instance, in 2019, the mental health research output from Africa was only 2% of the total research output, which is significantly less than all other WHO regions (WHO, 2021).

Overall, there is a shortage of ASD research funding, a high number of undiagnosed autistic children and autistic individuals who do not receive support or education at the appropriate healthcare or educational facility respectively. Consequently, SA has little potential for centralised ASD data collection that would facilitate building large ASD cohorts such as those present in HICs. This negatively impacts autism research, in particular genetic studies that typically require large sample sizes. It is critical to address these obstacles because research in Africa has the potential to make unique contributions to autism research. One of the most significant research opportunities in SA is its rich genetic diversity. African populations have the highest levels of genetic variation on the globe. This, along with the reduced linkage disequilibrium, increases the probability of pinpointing causal loci that would be beneficial to autism research on a global scale (Sirugo *et al*., 2019). Thus, large-scale hypothesis-free studies (like GWAS and epigenome-wide screens) could capture a broad view of the unique genetic architecture of autism in SA which can be the catalyst for understanding molecular mechanisms contributing to autism (Sirugo *et al*., 2019). The latter then presents opportunities for the identification of novel treatment drug targets. Despite the barriers to molecular research, SA has the capacity to improve autism research and make valuable contributions to the understanding of underlying molecular mechanisms in ASD.

## CONCLUSION

In this review, we report a critically low number of molecular ASD publications from SA compared to the UK, the USA, Brazil, and India. The uniquely diverse gene pool of African populations provides a unique resource for contributing novel insights into ASD pathophysiology. Addressing the shortage of molecular research in SA and SSA is therefore paramount to the advancement of global autism research. As government and international funding are crucial for propelling research capacity in SA, funding priorities need to be shifted to redress the shortfall in molecular ASD research and grant SA autism researchers an enhanced presence in the global autism research space.

