## Supplementary Table 1 for "Molecular autism research in Africa: a scoping review comparing publication outputs to Brazil, India, the UK, and the USA"

### SUPPLEMENTARY INFORMATION

**Supplementary Table S1**: First author and year for each publication identified in each thematic area from the broad SA ASD search in this review

| Social and family | |
|---|---|
| *Author* | *Year* |
| Meiring | 2016 |
| Pottas | 2016 |
| Schlebusch | 2016 |
| Schlebusch | 2017 |
| Wetherston | 2017 |
| Van Der Merwe | 2017 |
| Cole | 2017 |
| Mthombeni | 2018 |
| Alhazmi | 2018 |
| Clasquin-Johnson | 2018 |
| Schlebusch | 2018 |
| Dawson-Squibb | 2019 |
| Reddy | 2019 |
| Ramseur | 2019 |
| Viljoen | 2019 |
| Makombe | 2019 |
| Leigh | 2020 |
| Britz | 2020 |
| Simelane | 2020 |
| Dawson-Squibb | 2020 |
| Soeker | 2020 |
| Mazibuko | 2020 |
| Adams | 2020 |
| Adams | 2021 |
| Mupaku | 2021 |

| Clinical/ phenotyping/ behavioural | |
|---|---|
| *Author* | *Year* |
| Hoogenhout | 2016 |
| Hoogenhout | 2017 |
| Fourie | 2017 |
| Leclezio | 2018 |
| Naidoo | 2018 |
| Viviers | 2020 |
| Herdien | 2021 |
| Ringshaw | 2021 |
| Adams | 2021 |

| Education | |
|---|---|
| *Author* | *Year* |
| Hutton | 2016 |
| Mithimunye | 2018 |
| Erasmus | 2019 |
| Numisi | 2020 |
| de Jager | 2020 |
| Murray | 2020 |
| Sefotho | 2021 |
| Erasmus | 2021 |

| Prevalence/ demographics | |
|---|---|
| *Author* | *Year* |
| Rasdien | 2019 |
| Erasmus | 2019 |
| Pillay | 2021 |
| Pillai | 2021 |

| Professional knowledge | |
|---|---|
| *Author* | *Year* |
| Wium | 2018 |
| Krynauw | 2018 |

| Interventions/ treatment |
|---|

| *Author* | *Year* |
|---|---|
| Wallace | 2016 |
| Guler | 2018 |
| Hampton | 2019 |
| Franz | 2022 |
| Molecular | |
| *Author* | *Year* |
| O'Connell | 2018 |
| Stathopoulos | 2020 |
| Bam | 2021 |
| Naidoo | 2020 |
| Lee Evetts | 2021 |
| Diagnosis | |
| *Author* | *Year* |
| Smith | 2017 |
| Chambers | 2017 |
| Chambers | 2018 |
| Franz | 2018 |
| van Biljon | 2019 |
